# Homeostatic neuroimmune rhythms are linked to priming of olfactory bulb responses to an intranasal inflammatory challenge

**DOI:** 10.1101/2025.04.24.647555

**Authors:** Gregory Pearson, Brennan Baca, Jiexin Wang, Stephanie Gottwals, Nathan Santos, Giancarlo Denaroso, Amy Burnside, Saïd Akli, Ilia Karatsoreos

**Author notes:** To whom correspondence should be addressed: Ilia Karatsoreos, Ph.D., Department of Psychological and Brain Sciences 135 Hicks Way, Tobin Hall, University of Massachusetts Amherst Amherst MA, 01003, USA.

## Abstract

The circadian and immune systems are important for tissue homeostasis, yet their integration in the brain remains understudied. The olfactory bulb, a brain region that exhibits robust circadian rhythms and is regularly exposed to inflammatory stimuli, provides an optimal locus to probe the interaction of these two systems. We found that the murine olfactory bulb rhythmically expresses immune-related transcripts, with antiviral transcripts peaking around dusk. This was accompanied by distinct transcriptional responses to intranasal poly(I:C) at dusk versus dawn, suggesting that time of day primes the olfactory bulb’s response to inflammatory challenges. Using imaging flow cytometry, we detected two distinct populations of microglia, the resident macrophages of the brain, which differentially responded to intranasal poly(I:C) depending on time of day. This unveils a clear relationship between time of day and olfactory bulb immune processes, suggesting time is an important dimension to consider when studying the olfactory pathway into the brain.

## INTRODUCTION

The ability to predict recurring environmental changes or physiological processes enables organisms to survive and thrive in their environments. This ability is partly regulated by the circadian timing system, which enables cells, tissues, and organisms to anticipate daily environmental changes and to coordinate physiological processes throughout the day.^1^ In addition, organisms must also react to threats in their environment, such as injury or exposure to pathogens. The immune system is critical for responding to such threats, but it is becoming increasingly clear that it also plays an important role in tissue homeostasis.^2^

Both circadian and immune systems are important regulators of tissue homeostasis and play a significant role in health and disease. For example, the severity of infection is impacted by the time of day of pathogen exposure, and has been documented for various infection routes and pathogen tropisms and types.^3–9^ Additionally, circadian rhythm disruption influences both peripheral and brain inflammatory responses to immune challenges, and these altered responses are linked to prolonged sickness behaviors and sepsis severity^10–13^. In humans, chronic circadian disruption is associated with an increased risk of autoimmune conditions (e.g. psoriasis) and immune-related diseases (e.g. obesity, diabetes, cardiovascular disease, and cancer).^14–19^ Thus, circadian rhythms and immune function are linked. Yet how they are integrated, particularly in the brain, remains poorly understood.

The brain is an anatomically and functionally heterogeneous organ, ranging from sensory regions to highly integrative cortical regions. One such sensory region is the olfactory bulb, which processes the detection of odorants, enabling organisms to smell. Given its proximity and direct connection to the nasal cavity, the olfactory bulb is regularly exposed to environmental inflammatory stimuli.^20^ It also serves as a gateway for neurotropic pathogens, such as poliovirus, influenza A virus, and herpes simplex virus into the brain of humans, non-human primates, rodents, and other species^21,22^ The olfactory bulb also contains a molecular circadian clock that promotes daily changes in neuronal activity and odor discrimination.^23–26^ Intriguingly, time of day of intranasal exposure to a neurotropic virus significantly impacts the severity of infection.^5^ Taken together, circadian and immune systems are clearly implicated in olfactory bulb function. However, it remains unclear whether daily rhythms in immune processes occur in the olfactory bulb and how these underlying rhythms gate the response to a localized immune challenge.

Here, we demonstrate diurnal oscillations in the olfactory bulb’s immune state and investigate how these daily changes influence its response to an intranasal poly(I:C) challenge. We show that transcripts related to immune processes and function of microglia, the brain’s resident macrophages, are rhythmically expressed in the olfactory bulb under homeostatic (non-challenged) conditions. Furthermore, we find that rhythmic immune-related transcripts peaked around dark onset, suggesting a coordinated expression at this time. Finally, we illustrate that the type I interferon response to intranasal poly(I:C) occurs sooner following a challenge at dark onset compared to light onset, which is accompanied by a greater change in microglial state. Altogether, we propose that daily transcriptional changes in immune processes prime the olfactory bulb to differentially respond to an intranasal poly(I:C) challenge. Our findings implicate circadian rhythms as a key regulator of brain immune processes.

## RESULTS

### Neuroinflammation-related transcripts are rhythmically expressed in the olfactory bulb

Given that mediators of neuroinflammation are important for homeostatic brain functions,^27,28^ and that the olfactory bulb contains a molecular circadian clock,^23,24^ we hypothesized that the olfactory bulb will rhythmically express neuroinflammation-related transcripts under homeostatic conditions. To test this, we extracted RNA from olfactory bulbs of male mice collected every 3 hours (8 times of day) and used Nanostring nCounter technology to measure the expression of 757 neuroinflammation-related transcripts (Figure 1A). Using DiffCircaPipeline,^29^ we detected rhythmic expression of 224 neuroinflammation-related transcripts (30.5% of the 734 transcripts above background) (Figures 1B and 1C). Further, we validated our approach using qPCR to measure the expression of two core clock genes, *Per2* and *Nr1d1,* in our olfactory bulb samples. As expected, *Per2* and *Nr1d1* were rhythmically expressed according to DiffCircaPipeline analysis (Figures S1A and S1B). Together, these data demonstrate that the olfactory bulb rhythmically expresses neuroinflammation-related and clock-related transcripts.

**Figure 1.**
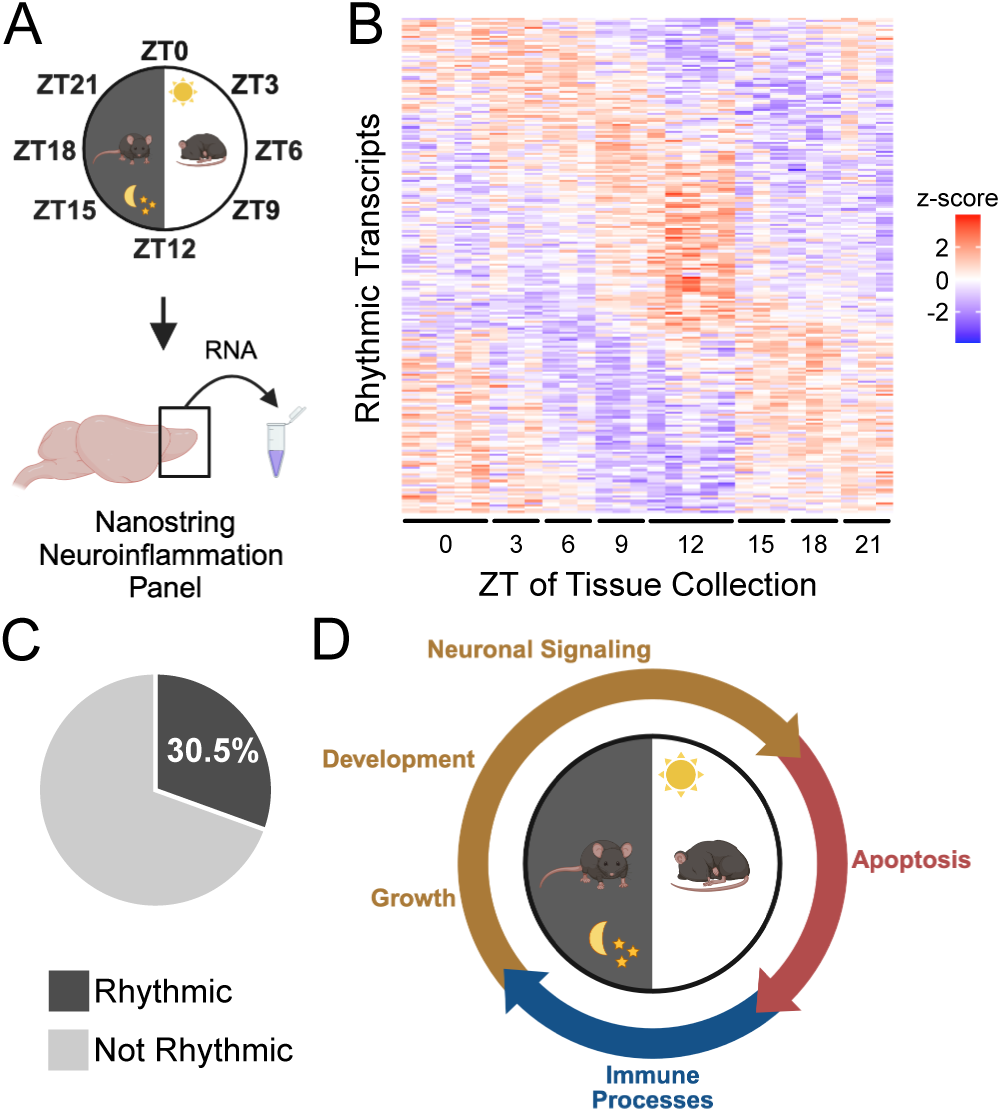
Neuroinflammation-related transcripts are rhythmically expressed in the olfactory bulb. (A) Schematic of experimental design. *n* = 3-5 mice per group. (B) Heatmap depicting rhythmic neuroinflammation transcripts. Normalized counts are represented as z-scores. (C) Pie chart showing the proportion of rhythmic and non-rhythmic neuroinflammation transcripts. (D) Summary diagram of Fig. S1C-S1F. Rhythmic neuroinflammation transcripts peaking within 6-hour phases of the day were thematically grouped into categories: growth, development, neuronal signaling, apoptosis, and immune processes. See also Figure S1 and Table S1.

We next investigated which biological processes these rhythmic neuroinflammation-related transcripts were involved in and whether they were coordinated in time. We organized the rhythmic transcripts into 6-hour bins based on when their rhythms peaked. The 6-hour bins included times around mid-dark (Zeitgeber Time (ZT); ZT15-ZT21), light onset (ZT21-ZT3), mid-light (ZT3-ZT9), and dark onset (ZT9-ZT15). We identified biological processes enriched within our 6-hour bins using Metascape.^30^ We further grouped them into four thematic categories based on our interpretation of these biological processes. These categories included growth and development, neuronal signaling, apoptosis, and immune processes (Figures 1D and S1C-S1F). We found that rhythmic transcripts related to growth and development peaked around mid-dark and light onset (Figures 1D, S1C, and S1D), while those related to neuronal signaling peaked primarily around light onset (Figures 1D and S1D). Additionally, transcripts related to apoptosis peaked around mid-light (Figures 1D and S1E). Notably, transcripts related to immune processes peaked around dark onset (Figures 1D and S1F), a time at which we have previously shown enhanced survival to a neurotropic virus infection.^5^ These data indicate that rhythmic neuroinflammation-related transcripts in the olfactory bulb are coordinated in time and that those peaking around dark onset are associated with immune processes.

### Antiviral transcripts are enriched at dark onset in the homeostatic olfactory bulb

Analysis of transcriptional rhythms in the olfactory bulb revealed a coordinated pattern of expression throughout the day (Figure 1D). Immune-related transcriptional rhythms peaked between ZT9 and ZT15 (Figure S1F), while other biological process-related transcriptional rhythms peaked between ZT15 and ZT9 (Figures S1C, S1D, and S1E). As such, we predicted that the olfactory bulb would have increased expression of immune transcripts at ZT12 compared to ZT0 (the midpoints of these time spans) under homeostatic conditions. To confirm this, we investigated the differential expression of neuroinflammation-related transcripts in the homeostatic olfactory bulb at these two times of day (Figure 2A). We found that neuroinflammation-related transcripts were primarily upregulated at ZT12 compared to ZT0 (Figure 2B). Using Metascape, we found that transcripts enriched at ZT12 were associated with antiviral defense and cytokine signaling pathways (Figures 2C, 2D, and S2A). Conversely, transcripts enriched at ZT0 were associated with cell junction assembly and TGF-β receptor signaling pathways (Figures 2C and S2B). Together, these data confirm that antiviral transcripts are enriched in the olfactory bulb at ZT12 and suggest that the olfactory bulb may be primed to more effectively respond to intranasal virus infections at this time of day.

**Figure 2.**
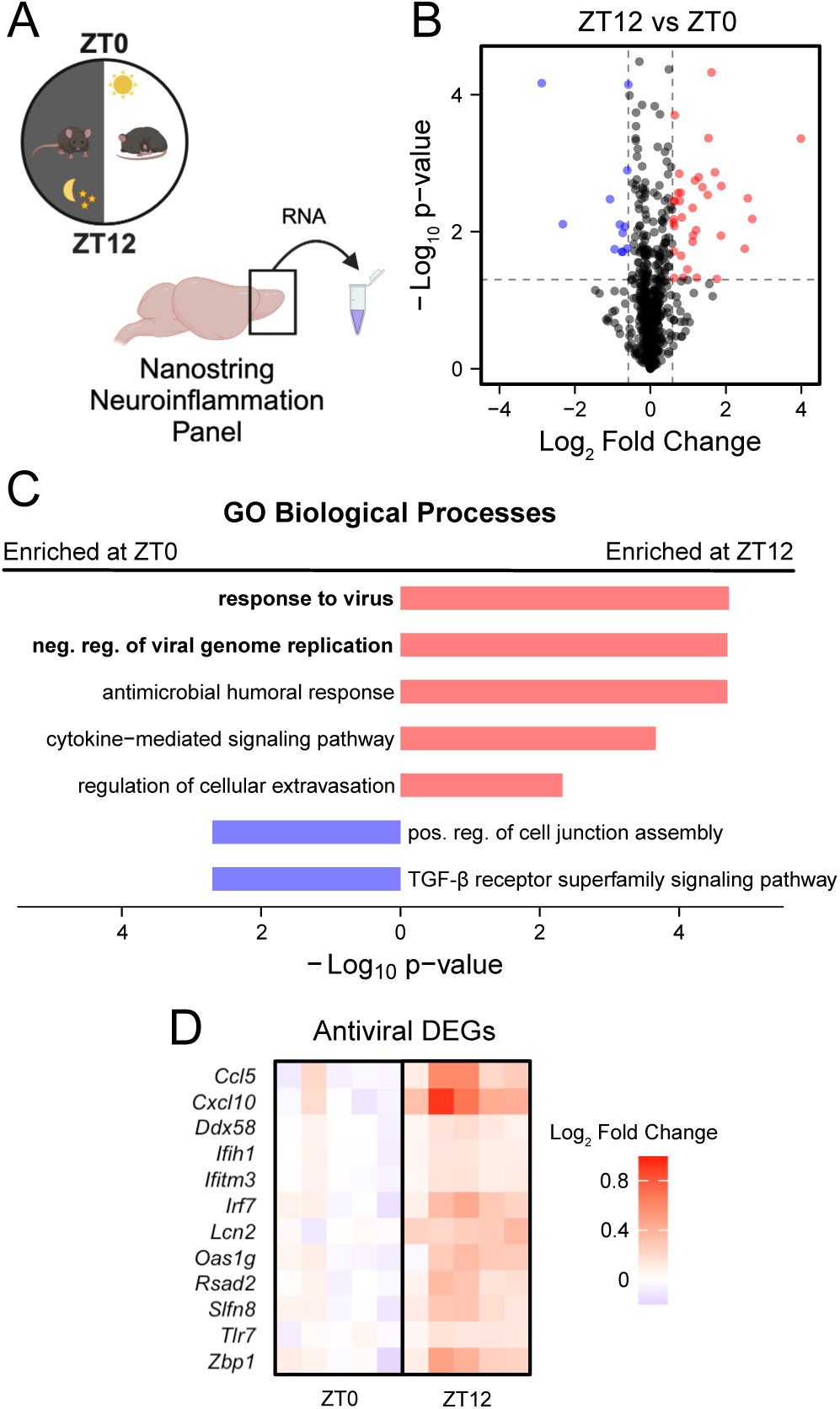
Antiviral transcripts are enriched at dark onset in the homeostatic olfactory bulb. (A) Schematic of experimental design. *n* = 5 mice per group. (B) Volcano plot showing differentially expressed genes (DEGs) that were enriched at ZT12 (red) or enriched at ZT0 (blue). Vertical dashed lines, log_2_ fold change = ±0.58. Horizontal dashed line, unadjusted *p* = 0.05. (C) Gene set enrichment analysis depicting biological processes enriched at ZT12 (red) and enriched at ZT0 (blue). Biological processes related to antiviral defense are bolded. (D) Heatmap depicting DEGs from the bolded biological processes related to antiviral defense from (C). Normalized counts are represented as log_2_ fold change relative to the mean of the ZT0 group. See also Figure S2.

### Time of day of intranasal poly(I:C) alters transcriptional response dynamics in the olfactory bulb

The olfactory bulb’s response to acute intranasal immune challenges is broad, including cytokine and chemokine production, microglial and astrocyte activation, and peripheral immune cell infiltration.^31,32^ Given that our differential expression analysis of the homeostatic olfactory bulb suggested priming of antiviral responses at ZT12 (Figure 2), we hypothesized that the transcriptional response to an intranasal immune challenge at ZT12 would unfold differently than the same challenge at ZT0. To test this, we challenged mice intranasally at ZT0 and ZT12 with vehicle or poly(I:C) (a synthetic analog of double-stranded viral RNA), collected tissues at 0-, 3-, 12-, and 24-hours post-inoculation, and assessed neuroinflammation-related transcriptional responses (Figure 3A). It is important to note that, to our knowledge, no published studies have investigated the response of the olfactory bulb to an acute intranasal poly(I:C) challenge. We found that time of day altered the olfactory bulb’s transcriptional response to an intranasal challenge, particularly at 3- and 12-hours post-inoculation (Figures 3B, 3C, S3A, and S3B). For instance, by 3- and 12-hours post-inoculation, intranasal challenge at ZT12 but not at ZT0 induced the expression of transcripts related to type I interferon signaling and antiviral responses (Figures 3D, 3E, S3A, and S3B). Conversely, by 12 hours post-inoculation, intranasal challenge at ZT0 reduced the expression of transcripts related to general immune responses (Figure S3A), a pattern not observed following challenge at ZT12. By 24 hours post-inoculation, intranasal challenge at ZT0 and ZT12 showed converging responses, with increased expression of transcripts related to type I interferon signaling and antiviral responses (Figures 3B, 3C, S3A, and S3B). These data indicate that time of day alters olfactory bulb response dynamics to intranasal poly(I:C), with an earlier antiviral response for mice challenged at ZT12 and an early downregulation of more general immune pathways for mice challenged at ZT0.

**Figure 3.**
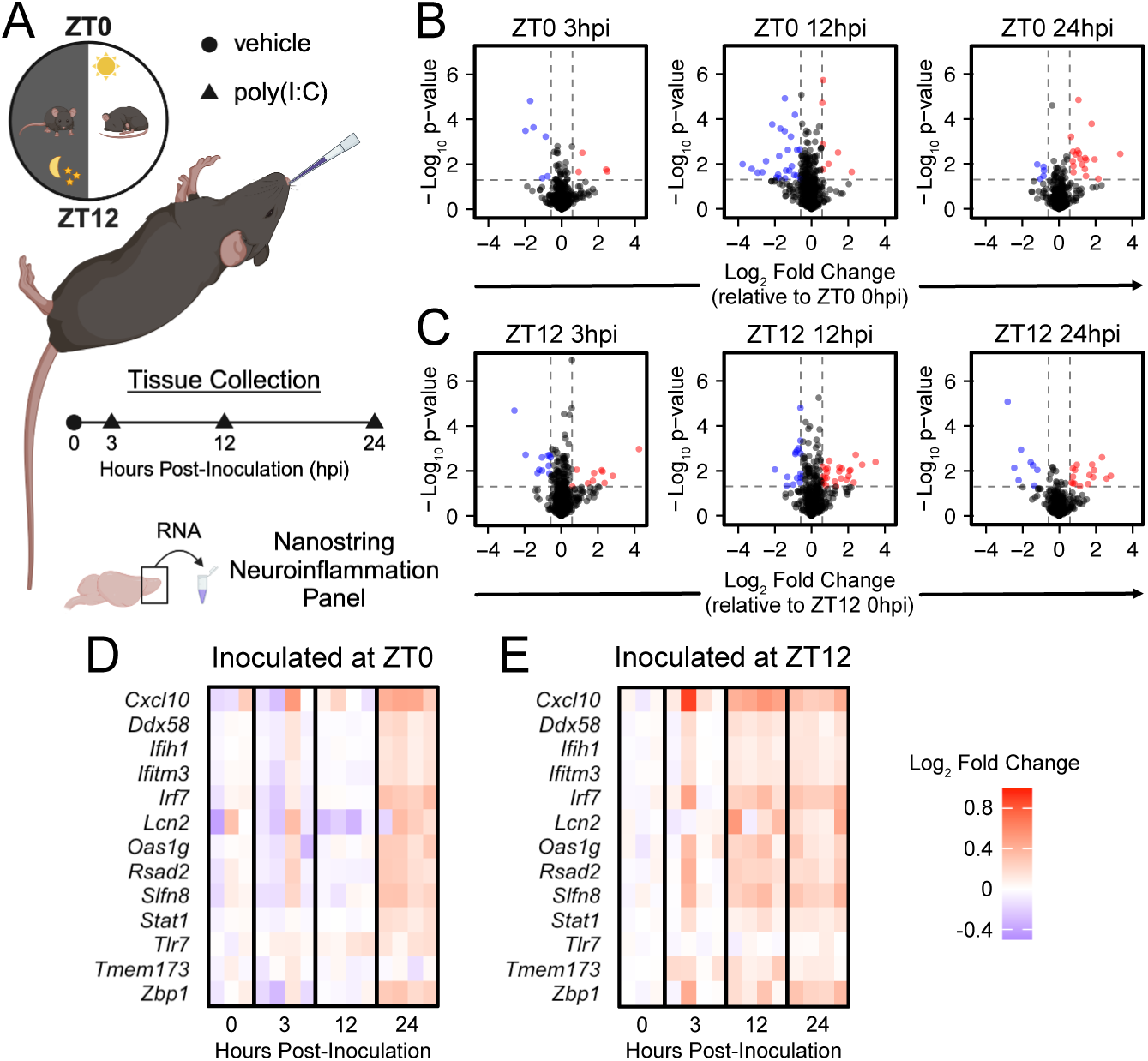
Time of day of intranasal poly(I:C) alters transcriptional response dynamics in the olfactory bulb. (A) Schematic of experimental design. *n* = 3-4 mice per group. (B, C) Volcano plots showing differentially expressed genes (DEGs) that were upregulated (red) or downregulated (blue) in response to poly(I:C) relative to vehicle control groups of the same ZT of inoculation. Vertical dashed lines, log_2_ fold change = ±0.58. Horizontal dashed lines, unadjusted *p* = 0.05. (D, E) Heatmaps depicting DEGs from the bolded biological processes related to antiviral defense and interferon signaling from (Fig. S3A-S3B). Normalized counts are represented as log_2_ fold change relative to the mean of the vehicle control groups. See also Figure S3.

### Microglial function related genes comprised the highest number of rhythmic transcripts in the homeostatic olfactory bulb

The olfactory bulb consists of multiple layers and numerous cell types.^33^ To identify cellular targets that might contribute to the differential responses to intranasal poly(I:C) we observed, we revisited our homeostatic rhythms data (Figure 1) and probed more deeply using nCounter’s pre-defined gene sets, which include neuron-, astrocyte-, oligodendrocyte-, and microglia-specific gene sets. We found that the microglia function gene set (which includes 187 transcripts) contained the greatest number of rhythmically expressed transcripts (*n* = 69, or 37%) (Figures 4A and 4B). This included 24 rhythmic transcripts specific to the microglia function gene set (Figure 4A). These data suggest that microglial functions in the olfactory bulb may change throughout the day.

**Figure 4.**
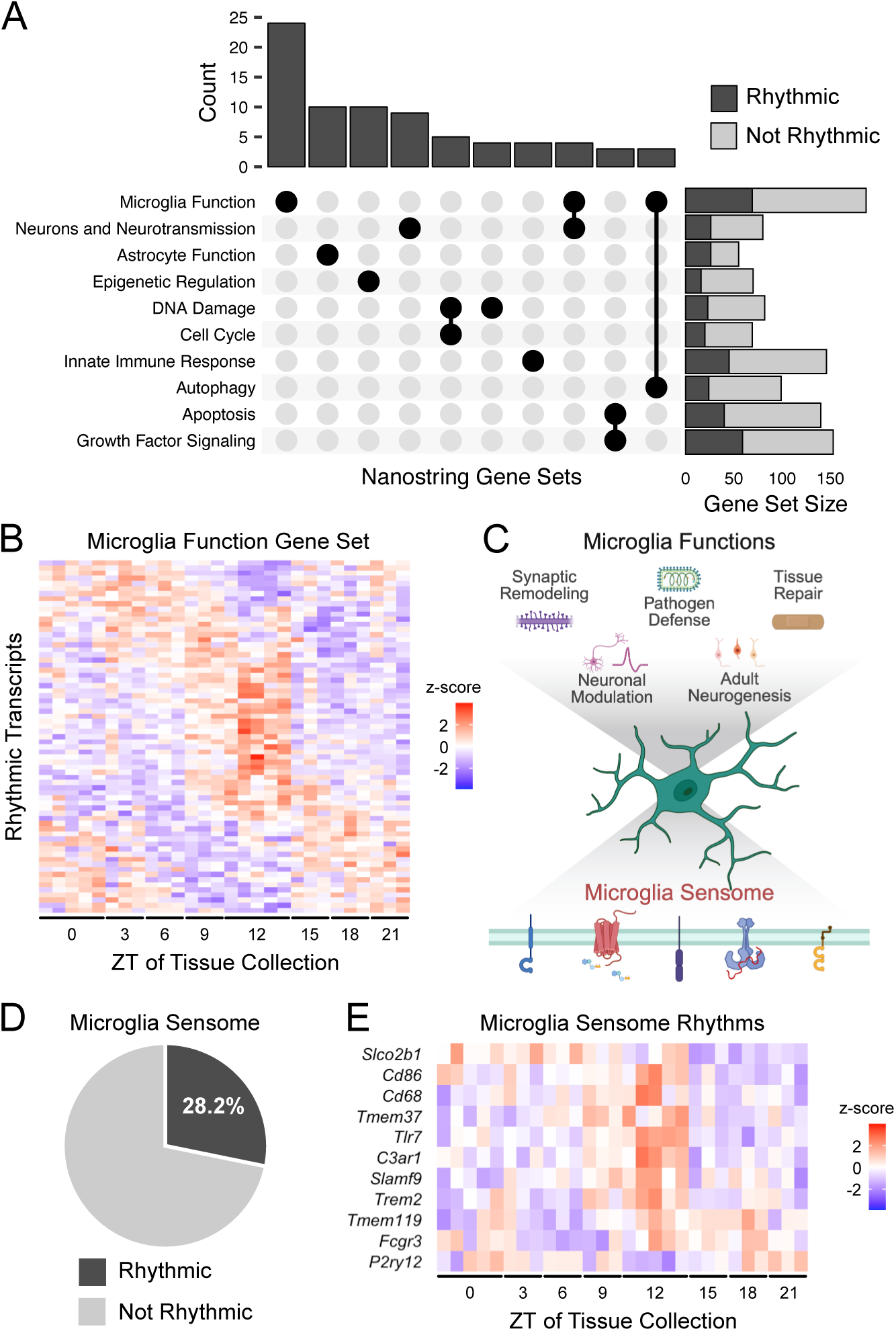
Microglial function related genes comprised the highest number of rhythmic transcripts in the homeostatic olfactory bulb. (A) UpSet plot showing the distribution of rhythmic neuroinflammation transcripts (from Fig. 1) into unique and overlapping Nanostring annotated gene sets. (B) Heatmap depicting rhythmic transcripts in the microglia function gene set. Normalized counts are represented as z-scores. (C) Graphical diagram depicting a selection of microglia functions, all of which rely on the microglia sensome to sense changes in their surrounding microenvironments. (D) Pie chart showing the proportion of rhythmic and non-rhythmic transcripts related to the microglia sensome. (E) Heatmap depicting rhythmic transcripts related to the microglia sensome. Normalized counts are represented as z-scores. See also Figure 1.

Microglia are involved in numerous brain functions and respond to abrupt changes in their surrounding microenvironment, including damage and inflammatory challenges (Figure 4C). This is accomplished collectively by a repertoire of microglial receptors termed the microglia sensome.^34^ Of the 757 neuroinflammation transcripts in the Nanostring nCounter panel, 39 were related to the microglia sensome, including 11 that were rhythmically expressed (28.2%) (Figures 4C-4E). These data suggest that time of day may influence olfactory bulb microglial responses to environmental perturbations.

### Olfactory bulb microglia consist of two distinct populations based on intrinsic fluorescence

Microglia are a heterogenous population that vary in morphological, molecular, and other cellular characteristics depending on various factors.^35^ To characterize olfactory bulb microglia, which make up approximately 8% of cells in the olfactory bulb,^36^ we used imaging flow cytometry to detect microglia based on the expression of surface proteins CD11b and CD45. We identified a population of cells characteristic of microglia (defined as CD11b^+^, CD45^low^) (Figures 5A, 5B, and 5C). This population consisted primarily of cells expressing the microglia-specific surface protein P2RY12 (Figures S4A and S4B). While acquiring these data, we observed intrinsically fluorescent (IF) spots within most olfactory bulb microglia (>80%) (Figures S5A and S5B). We defined IF as fluorescent emission detected in the 702/85 bandpass filter that was not attributed to spectral overlap from antibody-conjugated fluorophores (see Methods for more information). We also observed a bimodal distribution of IF intensity, indicating the presence of two distinct microglia populations, IF_low_ and IF_high_ (Figure 5D). While the majority (86%) of IF_low_ microglia did not contain IF spots, the majority (92.5%) of IF_high_ microglia contained one or more IF spots (Figure S5B). We found that these IF populations were also different in size (Figure 5E) and internal complexity (Figure 5F), with the IF_high_ population being larger and having more internal complexity than the IF_low_ population. Together, these data support the presence of two distinct populations of olfactory bulb microglia based on IF intensity. We propose that these populations represent different functional states and may be influenced by an inflammatory challenge.

**Figure 5.**
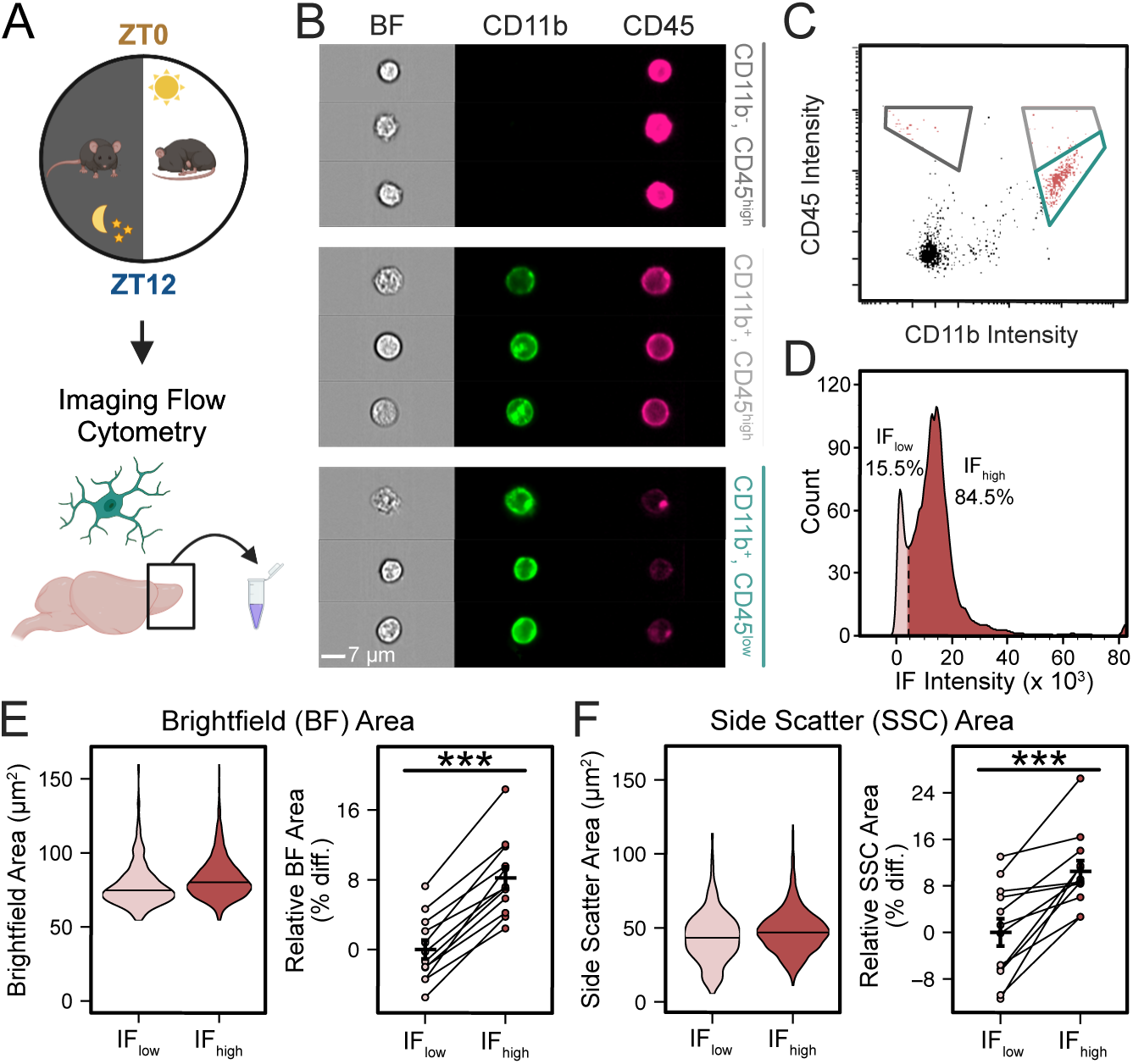
Olfactory bulb microglia consist of two distinct populations based on intrinsic fluorescence. (A) Schematic of experimental design. *n* = 6 mice per group. (B) Representative images of CD11b^-^, CD45^high^ (top, dark grey), CD11b^+^, CD45^high^ (middle, light grey), and CD11b^+^, CD45^low^ (bottom, turquoise) cells. Scale bar: 7 μm. (C) Representative gating strategy for differentiating CD11b^-^, CD45^high^ (dark grey gate), CD11b^+^, CD45^high^ (light grey gate), and CD11b^+^, CD45^low^ (turquoise gate) myeloid cell populations. (D) Histogram showing the bimodal distribution of olfactory bulb microglia (CD11b^+^, CD45^low^ cells) based on intrinsic fluorescence (IF) intensity. Data of microglia from all samples were pooled together to define the gate that distinguishes IF_low_ (light red) from IF_high_ (dark red) microglia. (E, F) Violin plots (left) and paired point plots (right) depicting differences between IF_low_ (light red) and IF_high_ (dark red) microglia in brightfield area (E) and side scatter area (F). The violin plots show the distribution of IF_low_ and IF_high_ microglia pooled together after data acquisition. The paired point plots show the median brightfield area (E) or side scatter area (F) for each biological replicate as a percent difference relative to the IF_low_ group. Error bars: mean ± SEM. Statistical significance, paired *t*-test: brightfield area, *t*_11_ = −11.9, *p* < 0.001; side scatter area, *t*_11_ = −5.34, *p* < 0.001. \**p* < 0.05, \*\**p* < 0.01, \*\*\**p* < 0.001. See also Figures S5 and S6.

### Intranasal poly(I:C) alters the proportion of olfactory bulb microglia populations in a time-of-day-dependent manner

Given that we observed the rhythmic expression of olfactory bulb transcripts related to the microglia sensome (Figure 4), we hypothesized that time of day would alter the response of olfactory bulb microglia to intranasal poly(I:C). To test this, we intranasally challenged mice with vehicle control or poly(I:C) at ZT0 or ZT12 and isolated microglia at 24 hours post-inoculation (Figure 6A). We detected a main effect of both ZT of inoculation and treatment on the proportion of IF_low_ and IF_high_ microglia populations in the olfactory bulb (Figure 6B). Notably, we did not detect an effect of time of day on the proportion of IF microglia populations under homeostatic conditions (Figures S6A and S6B), suggesting that the effect of time of day on IF microglia populations was dependent on an intranasal challenge. Intranasal poly(I:C) increased the proportion of IF_low_ microglia and decreased the proportion of IF_high_ microglia (Figure 6B), but more so at ZT12 than ZT0 (Figure 6C). These data indicate that time of day alters the response of olfactory bulb microglia to intranasal poly(I:C) and that microglial IF likely reflects cellular state and is impacted by environmental challenges.

**Figure 6.**
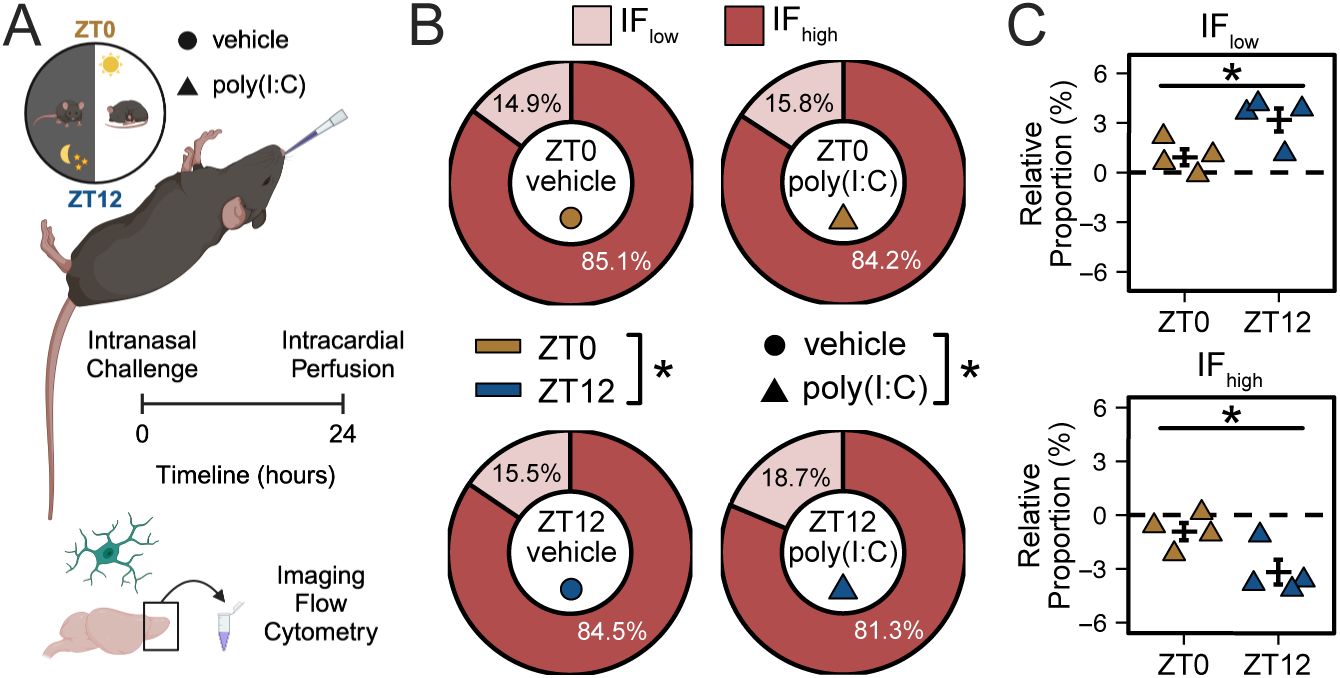
Intranasal poly(I:C) alters the proportion of olfactory bulb microglia populations in a time-of-day-dependent manner. (A) Schematic of experimental design. *n* = 4 mice per group. (B) Donut plots depicting the average proportion of IF_low_ (light red) and IF_high_ (dark red) microglia populations. Statistical significance, two-way ANOVA: intranasal treatment, *F*_1,12_ = 7.6, *p* = 0.017; inoculation ZT, *F*_1,12_ = 5.8, *p* = 0.033; interaction, *F*_1,12_ = 2.3, *p* = 0.15. \**p* < 0.05, \*\**p* < 0.01, \*\*\**p* < 0.001. (C) Dot plot showing the effect of intranasal poly(I:C) on the proportion of IF_low_ (top) and IF_high_ (bottom) relative to the mean of the vehicle controls from the same inoculation ZT. Error bars: mean ± SEM. Statistical significance, two-sample independent *t*-test: *t*_6_ = 2.67, *p* = 0.037. \**p* < 0.05, \*\**p* < 0.01, \*\*\**p* < 0.001.

## DISCUSSION

Our work provides several conceptual advances at the intersection of immunity and neuroscience. First, the olfactory bulb, a primary site of pathogen entry into the brain, displays remarkable daily rhythms in immune transcripts, even under homeostatic conditions. A large cohort of these rhythmic transcripts are linked to the function of microglia. Second, intranasal challenge with poly(I:C) generates distinct time-of-day-dependent transcriptional responses in the olfactory bulb. Finally, intrinsic fluorescence of olfactory bulb microglia reflects cellular state and is influenced by the time of day of intranasal challenge. Together, these results indicate that circadian rhythms influence the transcriptional state of the olfactory bulb, priming its response to immune challenge, with microglia likely playing a central role in maintaining tissue homeostasis.

The olfactory bulb shares characteristics with many neocortical brain areas. For instance, it comprises primary projection neurons and regulatory interneurons in a laminar structure, with intricate local circuitries.^33^ These characteristics enable the olfactory bulb to perform its primary function of receiving olfactory cues from the olfactory epithelium and transmitting those signals to cortical areas.^33^ Olfactory functions, including odor responsivity and discrimination, are circadian regulated^25,26^ and likely associated with daily changes in synaptic structure and function. Immune mediators, such as microglia, are well-documented in regulating synaptic remodeling and plasticity in other sensory systems during development and adulthood.^37–41^ Whether the daily transcriptional patterns in immune processes we observed in the olfactory bulb are shared with other brain regions will need to be addressed in future studies.

Despite similarities to other brain regions, the olfactory bulb is uniquely situated near the nasal cavity, making it directly accessible to the external environment. Therefore, the olfactory bulb is regularly exposed to inflammatory particles and serves as an important entry site into the brain.^20,21^ We demonstrate that intranasal poly(I:C) challenge induces distinct time-of-day-dependent transcriptional responses in the olfactory bulb, with an earlier type I interferon response at ZT12 than ZT0. Several potential mechanisms are consistent with this outcome. First, poly(I:C) may directly target pattern recognition receptors in the olfactory bulb, activating downstream signaling pathways and promoting antiviral-like responses. Second, the olfactory bulb may indirectly respond to poly(I:C) via signaling from the olfactory epithelium. The olfactory epithelium expresses toll-like receptor 3 (TLR3),^42^ a key pattern recognition receptor activated by poly(I:C).^43^ Activation of TLR3 induces a type I interferon response,^43^ leading to the production and secretion of type I interferons. Type I interferons signal surrounding cells via the type I interferon receptor (IFNAR).^44^ We found that IFNAR transcripts (*Ifnar1* and *Ifnar2*) were rhythmically expressed in the olfactory bulb. Remarkably, the expression of several antiviral transcripts that we found to be increased earlier in the response when challenged at ZT12 are regulated by Stat1, a transcription factor downstream of IFNAR activation.^45^ Together, our results lead to a novel hypothesis that the olfactory bulb may be primed to respond to challenge at different times of day.

Viral pathogens that enter the brain via the olfactory pathway can be detected by various pattern recognition receptors. Indeed, we showed that the mRNA expression of the pattern recognition receptors, retinoic acid-inducible gene-I (RIG-I, *Ddx58*) and melanoma differentiation-associated gene 5 (MDA5, *Ifih1*) are rhythmically expressed and higher at ZT12 than ZT0 under homeostatic conditions. Given that activation of RIG-I and MDA5 induces a type I interferon response,^46^ and that type I interferon signaling in the olfactory bulb is critical for the defense against neurotropic virus infections via the olfactory pathway,^44^ these findings provide rationale for explaining how mice intranasally infected with a neurotropic virus infection have greater survival when infected at ZT12 than ZT0.^5^

Of the many cell types present in the olfactory bulb, we posited that microglia may play an important role in this priming. Microglia are important regulators of neuroinflammation and contribute significantly to brain injury and neurodegenerative diseases.^47,48^ They are also necessary for defending against brain infections, including those targeting the olfactory pathway.^49^ Specifically, depletion of microglia leads to increased severity of intranasal neurotropic virus infection.^50,51^ Intriguingly, severity of a neurotropic virus infection via the olfactory pathway depends on the time of day of infection.^5^ This suggests there may be links between microglial state, time of day, and the response to an intranasal immune challenge.

Microglial states are dynamic and have been classified based on changes in gene and protein expression, morphology, and other physical characteristics.^35^ We revealed an interesting relationship between time of day and intrinsic fluorescence (IF) detected in olfactory bulb microglia. The presence of an IF signal in the olfactory bulb has been previously reported.^52^ It is sometimes viewed as an artifact limiting the ability to use fluorescence-based technology to obtain biological measurements, including of microglia.^53^ Recent work has suggested that IF is related to changes in lysosomal function, particularly during aging, and that some of it might be attributed to lipofuscin granules.^53,54^ Our work significantly expands the notion that IF is related to the current functional cellular state of microglia in the olfactory bulb. Specifically, our results demonstrate acute intranasal challenge with poly(I:C) not only quickly (within 24 hours) alters the proportion of IF_low_ and IF_high_ microglia populations in the olfactory bulb, but that it does so in a time-of-day-dependent manner, with a greater effect at ZT12 than ZT0. It is possible that this highly coordinated and temporally regulated response is, in part, enabled by the transcriptional rhythms of the microglia sensome.

Our data show that many aspects of olfactory bulb immune function are rhythmically coordinated on the circadian time scale. This raises the question of whether these rhythms impart any benefit? One possibility is that there is an evolutionary pressure to anticipate exposures to pathogens and mount more rapid responses at one time of day than another.^55^ Mice are nocturnal and highly olfactory-oriented. Thus, it is convenient to assume that when they are most active, they are more likely to be exposed to intranasal pathogens. Immune defense against pathogens should therefore be most protective during the active phase. In support of this, mice intranasally infected with a neurotropic virus at the onset of their active phase (ZT12, dark onset) have an approximately 50% higher survival than those infected at the onset of their resting phase (ZT0, light onset).^5^ However, when mice are intranasally infected with a non-neurotropic virus, like influenza virus, they have greater survival rates when infected near resting phase onset than near active phase onset.^9^ We propose these opposing outcomes argue against global circadian rhythms in immune defense. Instead, they suggest that the severity of infection depends not only on pathogen type, dose, and virulence factors, but also on the time of day of infection, as well as the tissue state prior to infection. As such, we posit that enhanced survival following a neurotropic virus infection at dark onset is a consequence of daily immune rhythms underlying olfactory bulb function, rather than an anticipation of a possible infection. That is, homeostatic rhythms in olfactory bulb functions prime the olfactory bulb to differentially respond to an intranasal immune challenge depending on time of day.

The findings we present represent two intersecting stories. The first is related to the classic role of the immune system in detecting, limiting the spread of, and clearing pathogens. The second is a less discussed role of the immune system: regulating tissue state and function.^2^ Our data support the overarching concept that both immune defense against pathogens and immune regulation of tissue function are dependent on time of day. Considering our findings and the known presence of an intrinsic molecular clock in microglia,^10,56^ we contend that microglia are positioned as key mediators of the circadian modulation of both aspects of immune function in the brain. Of course, additional studies are certainly warranted to fully establish this role for microglia. However, our results set the stage for several new lines of inquiry and strongly suggest the need to incorporate time of day into investigations of immune regulation of brain function. Indeed, we believe our findings have significant implications for how we consider immune processes in the brain: they are important for normal tissue function, not just infection or injury; and they are strongly influenced by time of day. Our work highlights a potential link between disrupted immune processes and altered tissue function, providing new insights into the etiology of neurodegenerative diseases, as circadian disruption is linked to Alzheimer’s, Parkinson’s, and other conditions.^57^ Our findings also have therapeutic implications, as the efficacy of nose-to-brain drug delivery^58^ may be significantly affected by time of day. In sum, our findings demonstrate that time is a critical dimension for understanding tissue function and how it relates to health and disease.

## LIMITATIONS OF THE STUDY

Our main interest in this study was to determine how time of day influences the immune profile of the brain, and if these daily changes are functionally relevant. We chose the olfactory bulb as our model tissue and investigated if these daily changes gate responses to an intranasal inflammatory challenge. The results we present provide a fascinating look into the temporal landscape of tissue level immune function of this brain region, opening several avenues of research. Future studies should probe the role of cell-autonomous endogenous rhythmicity in the olfactory bulb. Given it is well-documented that the olfactory bulb contains a robust circadian oscillator, determining if our findings hold in constant conditions and if they are linked to specific clock genes would be a logical next step. Our transcriptional data were derived from bulk tissue RNA and did not elucidate the contributions of different cell types or layers in the olfactory bulb. Using our foundational findings as a starting point, future studies using spatial transcriptomics and/or single-cell RNA sequencing will provide more clarity. Additionally, our studies were completed only in male mice due to technical and financial restrictions, and incorporation of female mice might provide additional insights. While our characterization of the IF signal in olfactory bulb microglia allowed identification of two microglial populations, the presence of IF makes it inherently difficult to use quantification methods dependent upon fluorescent signals. Application of techniques such as spectral flow cytometry or mass cytometry may assist in evaluating additional microglial characteristics. Finally, using poly(I:C) enabled us to focus primarily on TLR3 mediated responses. This work sets the stage to better understand the mechanisms by which circadian rhythms influence the outcome following exposure to pathogens via the nasal route. Such mechanisms might extend beyond direct impacts of time of day on the olfactory bulb; they will also likely provide insights on communication from the olfactory epithelium, as well as downstream signaling to other brain targets. Our foundational work was a necessary first step that would enable follow-up studies.

## METHODS

### Animals

Adult male C57BL/6N mice were received from Charles River Laboratories at 7-8 weeks old and housed in standard shoebox cages on a 12-hour light, 12-hour dark cycle with *ad libitum* access to food and water. Zeitgeber time 0 (ZT0) was defined as lights on, while ZT12 was defined as lights off. Cages were stored in sound-attenuated and ventilated isolation cabinets (Phenome Technologies, Skokie, IL) with a light intensity of 35-40 lux at cage level using broad spectrum white LEDs. Cabinet lighting schedules were controlled using ClockLab Chamber Control software (Actimetrics, Lafayette, IN). This unique design allowed us to perform mouse experiments from different ZT groups at the same facility time, controlling for potential extraneous variables associated with experimental manipulations. Room temperature was maintained at 21-23°C. Mice were 11-15 weeks old at the time of experiments/tissue collection. For homeostatic (i.e. non-challenged) gene expression experiments, mice were housed 3-5 per cage with a few exceptions (i.e. two ZT3 mice and three ZT6 mice were single-housed due to fighting). For all other experiments, mice were housed 1-2 per cage. All animal procedures were approved by the University of Massachusetts Amherst Animal Care and Use Committee.

### Intranasal Poly(I:C) Challenge

Mice were intranasally inoculated with vehicle or poly(I:C) (10 µL per nare, 2 mg/mL, polyinosinic-polycytidylic acid, high molecular weight, InvivoGen, Cat. Code: tlrl-pic) at ZT0 or ZT12. Even though mice were inoculated at one of two times of day, this always occurred in the light (i.e. immediately after lights on for ZT0 and immediately prior to lights off for ZT12) to avoid potential light pulse confounds. Poly(I:C) was dissolved in endotoxin-free physiological (0.9%) saline at 65°C for 10 minutes and stored at –20°C until experiments. We used 2 mg/mL of poly(I:C) since acute administration of this dose promotes IRF-3 phosphorylation and immune cell infiltration in the olfactory mucosa.^42^ For the gene expression experiment, mice were euthanized at 0-(vehicle treatment), 3-, 12-, or 24-hours post-inoculation via cervical dislocation. Brains were dissected and flash frozen in powdered dry ice. Brains were stored at −80°C until olfactory bulb removal and RNA extractions. For the imaging flow cytometry experiment, mice were transcardially perfused at 24-hours post-inoculation and olfactory bulbs were dissected to isolate microglia (see Microglia Isolations for Imaging Flow Cytometry for more information).

### RNA Extraction

Following removal from −80°C, olfactory bulbs were dissected on an ice-cold glass petri dish underlaid by crushed dry ice, collected into a tube, and stored at −80°C until RNA extractions. Olfactory bulbs were disrupted in the TissueLyser LT by rapid agitation (2 minutes at 50 Hz) in the presence of beads (5 mm steel) and QIAzol lysis reagent (Cat. # 79306). RNA was extracted using the RNeasy mini kit (Qiagen, Cat. # 74104) following the manufacturer’s instructions. Extracted RNA concentration/quality was analyzed using Qubit and NanoDrop prior to sending to UMass Boston Genomics Core for Nanostring analysis or cDNA synthesis for qPCR analysis.

### Nanostring Pre-processing

Transcriptional analysis of RNA samples extracted from olfactory bulbs was conducted using Nanostring nCounter technology. nCounter Murine Neuroinflammation panels allowed us to measure the impact of time of day and intranasal poly(I:C) challenge on the expression of 757 neuroinflammation-related mRNA transcripts. The nCounter panels were run at the CPCT Genomics Core at UMass Boston according to supplier protocols. Raw data were processed using Nanostring’s nSolver software (Version 4.0.70).

Quality control metrics were assessed to confirm quality of the data prior to data processing and are as follows: % Fields of View = 96-100%, binding density = 0.5-1.3, and positive control linearity = 0.98-1.00. All samples were within the manufacturer’s recommended range of acceptable quality control values.

Where batch-to-batch calibration was necessary, two calibrator samples were run twice, once on each batch. Calibrator samples were selected according to which samples had the greatest number of transcripts with raw counts above 100 as instructed by the manufacturer. Using these samples, data were batch calibrated using nSolver software. As recommended by the manufacturer, neither background subtraction nor background thresholding were performed. Genes within the panel that fell below background (i.e. had raw counts below 20 for 50% or more of the samples) were removed from the analysis.

### Nanostring Normalization

Data were normalized in a two-step process using both positive control and housekeeping normalization factors. The raw counts of each gene are multiplied by their lane-specific positive control normalization factor and then their lane-specific housekeeping normalization factor. The positive control normalization factors for each sample were within the recommended range of 0.3 to 3. Housekeeping genes used for calculating the housekeeping normalization factor were selected using the geNorm algorithm within the NormqPCR R library, which is incorporated within nSolver’s Advanced Analysis software (Version 2.0.134). This selects the optimal number of housekeeping genes using an iterative process of removing the most unstable housekeeping genes. For homeostatic gene expression analysis, the geNorm selected housekeeping probes were *Aars, Ccdc127, Cnot10, Csnk2a2, Fam104a, Lars, Mto1, Supt7l, Tada2b,* and *Xpnpep1*. For intranasal challenge gene expression analysis, all housekeeping probes used in the homeostatic gene expression analysis were used except for *Csnk2a2* as it was deemed unstable by geNorm. The housekeeping normalization factor was within the recommended range of 0.1 to 10 (10-fold range).

### Nanostring Rhythms and Differential Expression Analysis

Normalized data were then analyzed using DiffCircaPipeline^29^ to identify rhythmically expressed transcripts or nSolver’s Advanced Analysis to visually inspect the data, identify potential outliers, and perform differential expression analysis. For differential expression analysis in Advanced Analysis, the fast/recommended option was selected. The fast/recommended option infers differential expression using a negative binomial model. If the negative binomial model fails to produce a stable estimate of the parameters, then the fast/recommended option uses a log-linear regression to infer differential expression. Differentially expressed genes (DEGs) were defined as transcripts with an unadjusted p-value < 0.05 and a log_2_ fold change cutoffs of ≤ −0.58 or ≥ 0.58. A log_2_ fold change of 0.58 is equivalent to a fold change of 1.5 and we chose a 1.5-fold change to set a more conservative significance threshold. Rhythmic transcripts were defined as transcripts with a p-value < 0.05 following DiffCircaPipeline analysis. DiffCircaPipeline uses cosinor modeling to define rhythmicity parameters including phase, amplitude, Midline Estimating Statistic of Rhythm, and rhythm fitness.

### Nanostring Pathway Enrichment Analysis

Metascape^30^ was used to identify biological processes enriched in rhythmic transcripts or DEGs identified in our studies. The list of rhythmic transcripts or DEGs was added into Metascape and custom enrichment analysis for mouse was performed. The default settings for pathway and process enrichment were used with minimum overlap of 3, p-value cutoff of 0.01, and minimum enrichment of 1.5. The background genes were adjusted to include only the transcripts within the nCounter neuroinflammation panel that were above background. The GO Biological Processes database was selected for pathway enrichment analysis.

### Quantitative PCR

RNA extracted from olfactory bulbs was converted into cDNA using the High-Capacity RNA-to-cDNA^TM^ Kit (Applied Biosystems, Cat. # 4387406). Real-time qPCR was completed using iTaq^TM^ Universal Probes Supermix (Bio-Rad, Cat. # 1725131) and off-the-shelf TaqMan Gene Expression Assays (ThermoFisher), including *Rn18s* (Mm03928990_g1), *Per2* (Mm00478099_m1), and *Nr1d1* (Mm00520708_m1). Samples were run in triplicate using a 20 µL reaction volume on 96-well plates, which were run on a Bio-RAD CFX96 Touch Real-Time PCR Detection System. CT values were produced using Bio-Rad CFX Manager software (Version 3.1.1517.0823). Rn18s was used as the housekeeping gene. Fold change was calculated using the ΔΔCT method of relative quantification. All data were normalized to the group that was collected at ZT0. Fold change values were used for DiffCircaPipeline analysis.

### Microglia Isolations for Imaging Flow Cytometry

Mice were anesthetized using a ketamine (45 mg/mL)/xylazine (3 mg/mL) solution administered intraperitoneally. Following transcardial perfusion with ice-cold 1X PBS (Gibco, Cat. # 10010031), brains were dissected and briefly stored in ice-cold DMEM/F12 (Gibco, Cat. # 21041025) until all brains were collected. Olfactory bulbs were dissected and chopped with razor blades on an ice-cold glass petri dish and then enzymatically/physically dissociated and filtered to obtain single-cell suspensions as described by Moseman et al.^50^ Briefly, tissues were enzymatically digested in Collagenase D (Roche, Cat. # 11088858001, 1 mg/mL) and DNase I (Roche, Cat. # 10104159001, 0.25 mg/mL) dissolved in RPMI 1640 Medium (Gibco, Cat. # 11835030) for 30 minutes at 37°C. Enzymatic digestion was coupled with physical digestion by triturating every 10 minutes. After dissociation, samples were washed with ice-cold RPMI and centrifuged at 1500 x g for 3 minutes at 4°C. Next, myelin was removed from the single-cell suspensions by resuspending the samples in a 37% isotonic Percoll solution (Percoll, Sigma, Cat. # P4937; 10X PBS, Gibco, Cat. # 70011044; 1X PBS, Gibco, Cat. # 10010031) and centrifuged at 1500 x g for 15 minutes at 4°C (max acceleration, no brake). The myelin layer and Percoll supernatant were then removed through vacuum suction and the cell pellets were washed with ice-cold FACS buffer (2% FBS, Cytiva, Cat. # SH30071.03; 1 mM EDTA, Invitrogen, Cat. # AM9262; 1X PBS, Gibco, Cat. # 10010031) with centrifugation at 1500 x g for 3 minutes at 4°C. Single-cell suspensions were resuspended with ice-cold FACS buffer and transferred to a 96-well v-bottom plate for flow cytometry staining.

### Flow Cytometry Staining

After transfer to 96-well v-bottom plate, samples were washed with ice-cold PBS by centrifugation at 1500 x g for 3 minutes at 4°C and then stained with LIVE/DEAD Fixable Dead Cell Stain (Invitrogen, LIVE/DEAD Fixable Violet, Cat. # L34963) according to the manufacturer’s instructions. Briefly, samples were resuspended in the LIVE/DEAD Fixable stain (1:1000) and incubated on ice for 30 minutes and protected from light. Samples were then washed with PBS by centrifugation at 1500 x g for 3 minutes at 4°C. Following live/dead staining, samples were blocked in FACS buffer containing 0.0025 mg/mL of Fc receptor blocking antibody (TruStain FcX PLUS, BioLegend, Cat. # 156604, RRID: AB_2783137) for 10 minutes. After Fc receptor blocking, samples were stained in FACS buffer containing fluorophore-conjugated antibodies specific to extracellular targets (CD11b-BV510, 1:100, BD Biosciences, Cat. # 562950, RRID: AB_2737913; CD45-APC/Cy7, 1:200, BioLegend, Cat. # 103116, RRID: AB_312980; P2RY12-PE, 1:100, BioLegend, Cat. # 848003, RRID: AB_2721644) and incubated for 20 minutes on ice and protected from light. Cell suspensions were washed with FACS buffer, resuspended in FACS buffer, and transferred to 1.5 mL microcentrifuge tubes to be run on Amnis ImageStream MKII (Luminex). Acquired data were compensated, gated, and analyzed using Amnis IDEAS software (Version 6.2.187.0). Gating figures were created using both Amnis IDEAS and FlowJo (Version 10.10.0) software. Compensation was performed using UltraComp eBeads (Invitrogen, Cat. # 01-2222-41) for extracellular antibodies and Arc Amine Reactive Compensation Bead Kit (Invitrogen, Cat. # A10628) for LIVE/DEAD Fixable stain as per manufacturer’s instructions. Pooled samples were used as fluorescence minus one (FMO) controls. FMO controls were used to differentiate P2RY12 antibody signal from intrinsic fluorescence (IF) signal^59^ and it was determined that 89.3% ± 0.9% (mean ± SEM) of the CD11b^+^, CD45^low^ population expressed P2RY12 (Figure S4), confirming the analyzed population consisted of primarily microglia. We defined IF as fluorescent emission detected in the 702/85 bandpass filter. While IF appeared in all emission bandpass filters, the 702/85 filter was selected since IF signal in microglia exhibits maximal intensity at this wavelength.^54^ Additionally, IF was not attributed to spectral overlap from antibody-conjugated fluorophores as compensation was performed.

### Statistical Analysis

In addition to DiffCircaPipeline and DEG analyses described above, the following statistical analyses were performed. Paired and unpaired two-tailed *t*-tests were used to detect differences between two groups where appropriate. Assumptions of normal distribution and homogeneity of variance were tested using the Shapiro-Wilk Normality and Levene’s tests, respectively. Two-sample Wilcoxon rank sum exact test was used when comparing two groups with non-normal distributions. To test the effects of time of day and intranasal treatment on the proportion of olfactory bulb microglia populations, we performed two-way ANOVAs. Specific statistical analyses and results are included within figure captions. All analyses and figures were completed using R (Version 4.3.1).^60^

## ACKNOWLEDGEMENTS

We would like to thank and acknowledge the following people and organizations for their contributions to our work: Michelle MacGray and the animal care staff at UMass Amherst for their dedicated assistance with animal care; Dr. Shaon Sengupta, Dr. Ryan Logan, and Dr. Santanu Bose for their valuable input on the manuscript; Patricia Reis, Dr. Matthew Hartsock, and Dr. Adolfo Cuadra for their recommended edits, comments, and proofreading of the manuscript; the scientific cores at UMass Amherst and UMass Boston for their assistance in data acquisition, including the UMass Amherst Flow Cytometry and Genomic Resource Laboratory Cores and the UMass Boston Genomics Core; the Keck Foundation and the National Institutes of Health (R01 DK119811; R01 AG082349) to IK for funding this work; and BioRender.com for the platform to develop our illustrative diagrams.

## AUTHOR CONTRIBUTIONS

Conceptualization, G.P. and I.K.; experimental design, G.P. and I.K.; performed experiments, G.P., B.B., J.W., S.G., N.S., G.D., S.A, and I.K.; data curation, G.P.; analyzed data, G.P. and A.B.; funding acquisition, I.K.; project administration, G.P., B.B., and I.K.; visualization, G.P.; writing – original draft, G.P. and I.K; writing – review and editing, G.P. and I.K. with input from all authors.

## DECLARATION OF INTERESTS

The authors declare no competing interests.

## Notes

### Competing Interest Statement

The authors have declared no competing interest.

