## Supplemental Figures S1 to S6 for "Homeostatic neuroimmune rhythms are linked to priming of olfactory bulb responses to an intranasal inflammatory challenge"

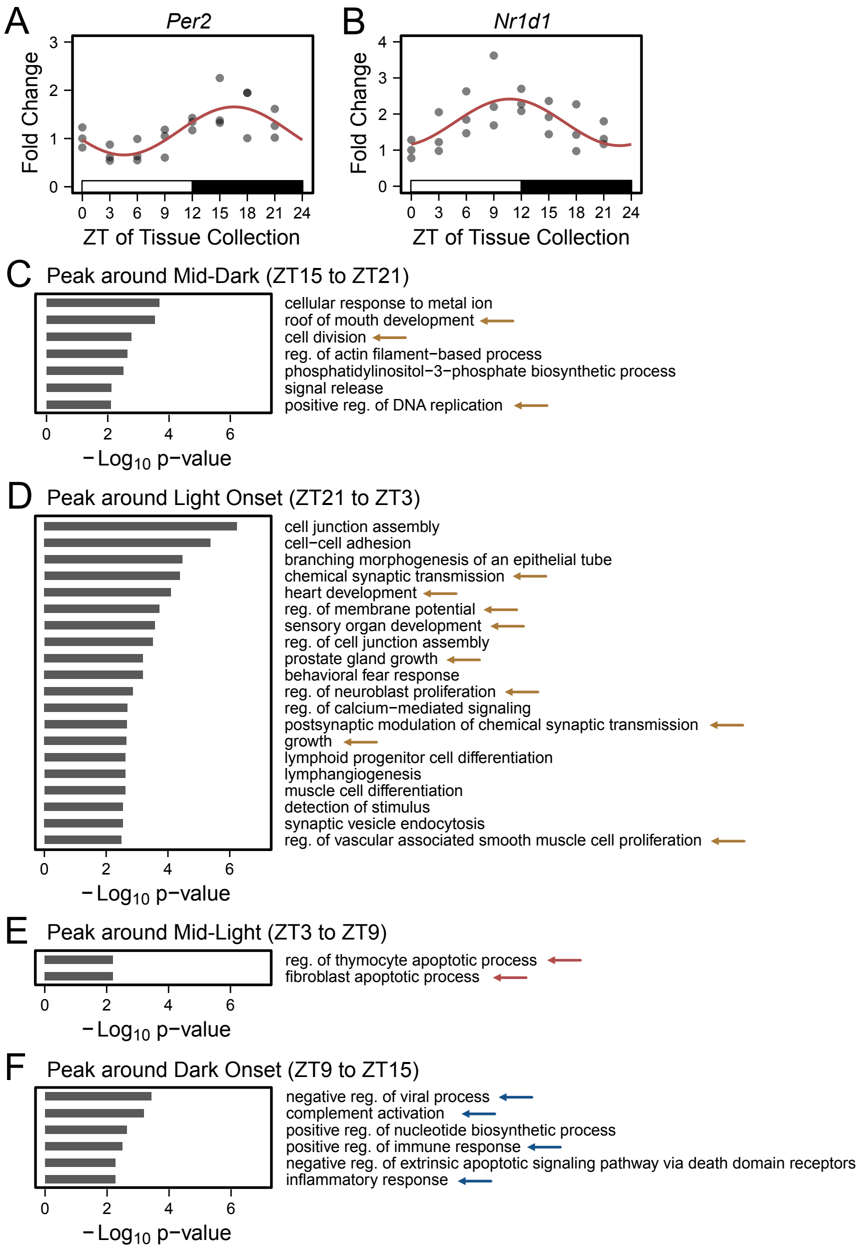


**Figure S1. The olfactory bulb rhythmically expresses core clock genes and neuroinflammation transcript rhythms are coordinated in time, Related to Figure 1.**

(A, B) Scatterplots with fitted cosinor curves to depict the rhythmic expression of the core clock genes *Per2* (A) and *Nr1d1* (B) in the olfactory bulb. *n* = 3 mice per group.

(C-F) Gene set enrichment analysis depicting pathways enriched in rhythmic transcripts peaking around mid-dark (ZT15-ZT21) (C), light onset (ZT21-ZT3) (D), mid-light (ZT3-ZT9) (E), and dark onset (ZT9-ZT15) (F). Arrows indicate biological processes related to growth, development, or neuronal signaling (yellow), apoptosis (red), or immune processes (blue) as highlighted in Fig. 1D.

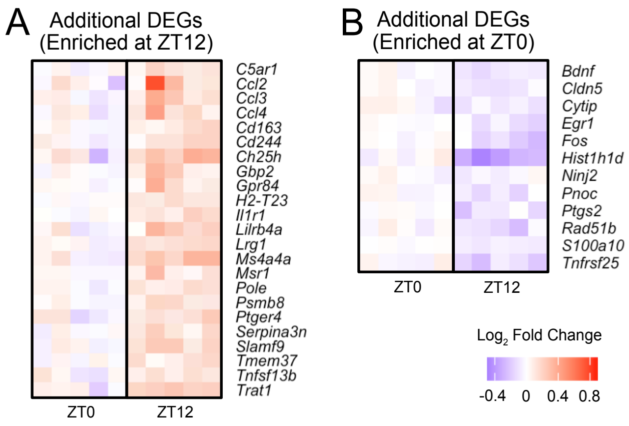


**Figure S2. Heatmaps of non-antiviral transcripts differentially expressed genes, Related to Figure 2.**

(A, B) Heatmaps showing additional differentially expressed genes (DEGs) enriched at ZT12 (A) or ZT0 (B). Additional DEGs are those DEGs that were not enriched in the antiviral-related pathways of Fig. 2C. Normalized counts are represented as $log_2$ fold change relative to the mean of the ZT0 group.

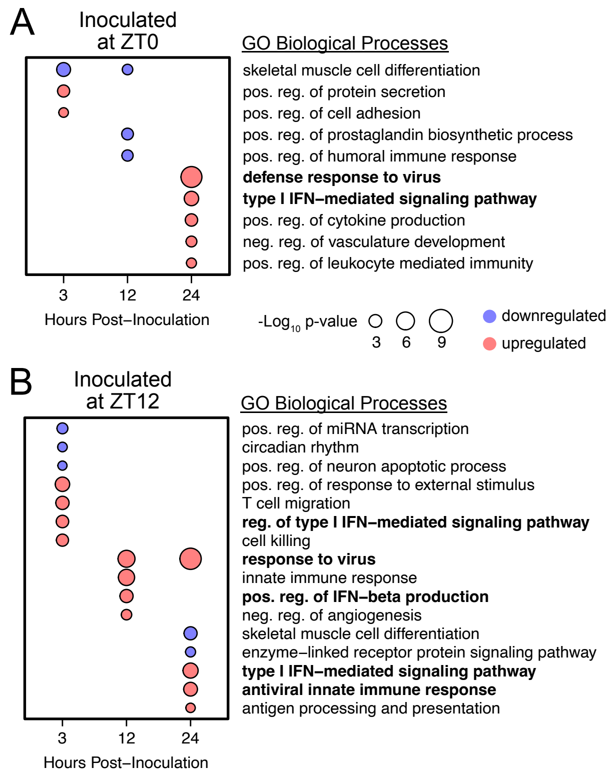


**Figure S3. Pathway analysis of differentially expressed genes in response to intranasal poly(I:C), Related to Figure 3.**

(A, B) Gene set enrichment analysis depicting biological processes of upregulated (red) or downregulated (blue) differentially expressed genes (DEGs) in response to intranasal poly(I:C) at ZT0 (A) or ZT12 (B). The size of the circles represents a change in $-Log_{10}$ p-value with larger sizes associated with greater significance. Biological processes related to antiviral defense or type I interferon signaling are bolded.

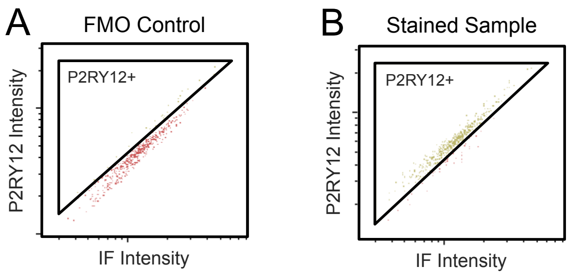


**Figure S4. Most of the analyzed olfactory bulb microglia population express P2RY12, Related to Methods.**

(A, B) Gating strategy for differentiating P2RY12+ (yellow) from P2RY12- (red) cells using an FMO control (A) and a representative stained sample (B). The vast majority of analyzed olfactory bulb microglia (CD11b$^{+}$, CD45$^{low}$ cells) expressed P2RY12, a microglia-specific purinergic receptor, confirming the population was enriched with microglia.

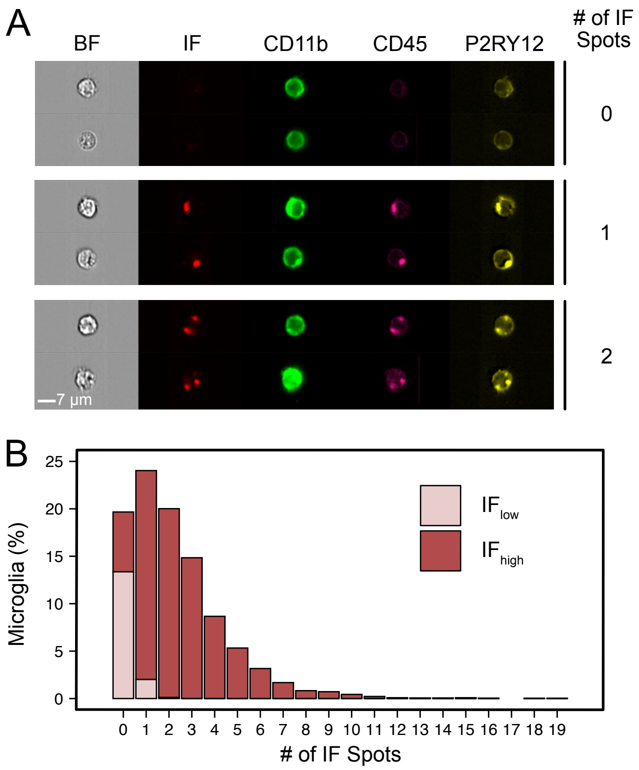


**Figure S5. Olfactory bulb microglia contain intrinsically fluorescent puncta, Related to Figure 5.**

(A) Representative images of $CD11b^{+}$, $CD45^{low}$ microglia with 0 (top), 1 (middle), or 2 (bottom) intrinsically fluorescent (IF) spots. The number of IF spots per microglia was determined using spot count analysis in the IDEAS software. Scale bar: 7 μm.

(B) Histogram showing the proportion of microglia containing differing numbers of spots. Data were pooled from all samples. The proportion of $IF_{low}$ (light red) and $IF_{high}$ (dark red) per spot count category was determined.

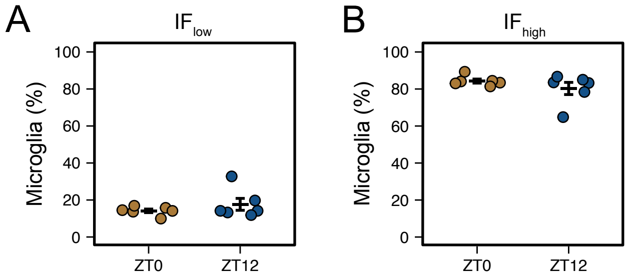


**Figure S6. Time of day does not influence the proportion of intrinsically fluorescent microglia population in the homeostatic olfactory bulb, Related to Figure 5.**

(A, B) Point plot showing the proportion of $IF_{low}$ (A) and $IF_{high}$ (B) microglia for each sample collected at ZT0 (yellow) or ZT12 (blue). Error bars: mean ± SEM. Statistical significance, two-sample Wilcoxon rank sum exact test: $W = 20$, $p = 0.82$.
